# Cartilage microRNA dysregulation in mouse osteoarthritis overlaps with patient disease candidates

**DOI:** 10.1101/113456

**Authors:** Louise H. W. Kung, Varshini Ravi, Lynn Rowley, Constanza Angelucci, Amanda J Fosang, Katrina M Bell, Christopher B Little, John F Bateman

**Author notes:** **Correspondence**: John F. Bateman, Murdoch Childrens Research Institute, Royal Children’s Hospital, Flemington Road, Parkville, Victoria, 3052, Australia.

## Abstract

To explore the role of microRNAs in osteoarthritis (OA), we conducted microRNA expression profiling on micro-dissected tibial cartilage and subchondral bone in a mouse model of OA produced by medial meniscus destabilization (DMM). DMM mice had characteristic cartilage degeneration, subchondral bone sclerosis and osteophyte formation. While subchondral bone showed no microRNA dysregulation, 139 microRNAs were differentially expressed in DMM cartilage at 1 and/or 6 weeks after OA initiation. To prioritize OA-candidates, dysregulated microRNAs with human orthologues were filtered using paired microRNA:mRNA expression analysis to identify those with corresponding changes in mRNA target transcripts in the DMM cartilage. An important cohort overlapped with microRNAs identified in human end-stage OA. Comparisons with microRNAs dysregulation in DMM mouse cartilage where aggrecan cleavage was genetically-ablated demonstrated that all were independent of aggrecan breakdown, earmarking these as important to the critical stages of OA initiation. Our comprehensive analyses identified high-priority microRNA candidates that have potential as human OA-biomarkers and therapeutic targets.

**SUMMARY:** Kung et al. conducted global analysis of microRNA dysregulation in joint tissues of a well-established mouse osteoarthritis model. Stringent filtering against human microRNA orthologues, integrated mRNA target analysis and comparison with published studies on human end-stage osteoarthritis identified microRNA candidates of potential clinical relevance.

## INTRODUCTION

Osteoarthritis (OA) is the most prevalent of all joint diseases, causing considerable morbidity, healthcare burden and financial expenditure (Cross et al., 2014; Puig-Junoy and Ruiz Zamora, 2015). Although many OA-related molecular pathways have been described, the key triggering molecular events remain elusive and as a result there are currently no registered therapies that halt disease onset or progression, only those which attempt to manage the associated pain (Hunter, 2011; Yu and Hunter, 2015). Thus, discovering key pathological pathways which may offer new, alternative targets for disease modifying therapies is a priority.

Articular cartilage degeneration is the characteristic feature of OA and it is accompanied by sclerosis of the underlying subchondral-bone (SCB), formation of osteophytes, ligament and meniscal damage along with synovial hyperplasia and inflammation, ultimately resulting in complete joint “organ” failure (Loeser et al., 2012). While cartilage degradation remains the hallmark of OA, it is clear that all joint tissues can contribute to the pathological process with the regulatory interplay between the cartilage and SCB thought to be of particular importance (Sharma et al., 2013; Yuan et al., 2014). However, the molecular characteristics of the disease pathways both within, and between these tissues remains poorly characterized, highlighting the critical need to perform parallel studies on molecular mechanisms on both articular cartilage and SCB during OA initiation and progression.

microRNAs (miRs) are an important class of cellular regulators which act by modulating gene expression at the post-transcriptional level during numerous physiological and disease settings (Kloosterman and Plasterk, 2006; Lewis and Steel, 2010; Sayed and Abdellatif, 2011). miRs are a family of evolutionarily conserved small non-coding RNAs, of which more than 2500 mature miRs have been identified in human to date (www.mirbase.org; (Kozomara and Griffiths-Jones, 2014)). Mature miRs range from 21-25 nucleotides in length and are formed from larger precursor miRs termed primary miRs (or pri-miR). The pri-miR are then processed by ribonuclease-III (RNase III) Drosha-DGCR8 micro-processing complex and subsequently Dicer to liberate mature miRs which then associate with Argonaute family proteins to form an RNA-induced silencing complex (RISC). These RISC effector complexes are then guided to target genes where base-pairing of the miR results in translational repression or mRNA decay of the target transcript (Ameres and Zamore, 2013; Ha and Kim, 2014; He and Hannon, 2004).

miRs have been shown to play critical roles in chondrocyte proliferation and differentiation during skeletal growth, as cartilage specific ablation of *Dicer* in mice resulted in skeletal abnormalities, decreased chondrocyte proliferation and accelerated differentiation into hypertrophic chondrocytes (Kobayashi et al., 2008), the latter of which is a key OA-response (Bateman et al., 2013; Little and Hunter, 2013a). Moreover, early postnatal deletion of *Drosha* in articular chondrocytes resulted in increased cell death and decreased matrix proteoglycan content (Kobayashi et al., 2015), both also key pathophysiological features of OA. These studies demonstrate the critical role of miRs globally in cartilage development, health and disease. An increasing number of specific miRs have been suggested as regulators of chondrocyte-driven processes central to OA pathology (*see Reviews* (Gibson and Asahara, 2013; Miyaki and Asahara, 2012; Nugent, 2016; Swingler et al., 2012; Trzeciak and Czarny-Ratajczak, 2014; Vicente et al., 2016)) making miRs viable new candidates for therapeutic targets and clinical biomarkers. Discovering their role in OA pathogenic processes is an exciting new frontier in unravelling the molecular mechanisms of OA to develop new mechanistically based therapies.

In this study we explore miR expression in articular cartilage and underlying SCB during the early stages of disease initiation and progression in a mouse model where OA was surgically-induced via destabilisation of the medial meniscus (DMM). This well-established, widely accepted mouse model of post-traumatic OA (which represents ~25% of all knee OA cases) not only informs on OA pathogenesis in general (Little and Hunter, 2013b; Malfait and Little, 2015) but also allows the molecular interrogation of the early stages of disease not possible with patient tissues. Studying these early stages of the disease is pivotal as it represents a critical time when therapeutic intervention is expected to have the most potential to delay disease progression and improve patient outcomes. We identified miR dysregulation specifically in OA cartilage and compared our data with previous human OA studies. *In silico* target prediction analysis on paired miR:mRNA data identified miR candidates with targets and functional biological processes relevant to human OA pathology. Furthermore, by comparing OA-related miR candidates in wild-type mice with their expression following DMM mice in which aggrecan breakdown was genetically ablated, we were able to discriminate between miRs which are independent of, or dependent on, aggrecan breakdown to provide additional insights into the sequence of miR dysregulation in the onset and progression of disease. Our study describes for the first time potential miR regulators of the initiation and progression of OA cartilage degeneration, not only validating those that have been shown in end-stage human OA but also identifying novel miRs not previously associated with OA, that may be potential therapeutic targets for this recalcitrant disease.

## RESULTS

### Histological characterization of articular cartilage and SCB of DMM and sham-operated mice

DMM surgery induced a post-traumatic OA pathology characterized by chondrocyte hypertrophy, progressive proteoglycan loss, structural damage and SCB sclerosis (Fig. 1) as previously reported (Jackson et al., 2014; Shu et al., 2016). Chondrocyte hypertrophy was increased at both 1 and 6 weeks (p = 0.002 and 0.004, respectively), but did not progress with time in DMM between 1 and 6 weeks (Fig. 1A). There was no change in proteoglycan loss in DMM compared with sham-operated joints at 1 week, but significant proteoglycan loss was observed in the DMM group with time (p = 0.0002) and in DMM versus sham at 6 weeks (p = 0.0002; Fig. 1B). Similarly, total cartilage structural damage scores did not differ between DMM and sham at 1 week, but scores increased with time in DMM (p = 0.01) and were greater in DMM versus sham at 6 weeks (p = 0.0002; Fig. 1C).

**Figure 1.**
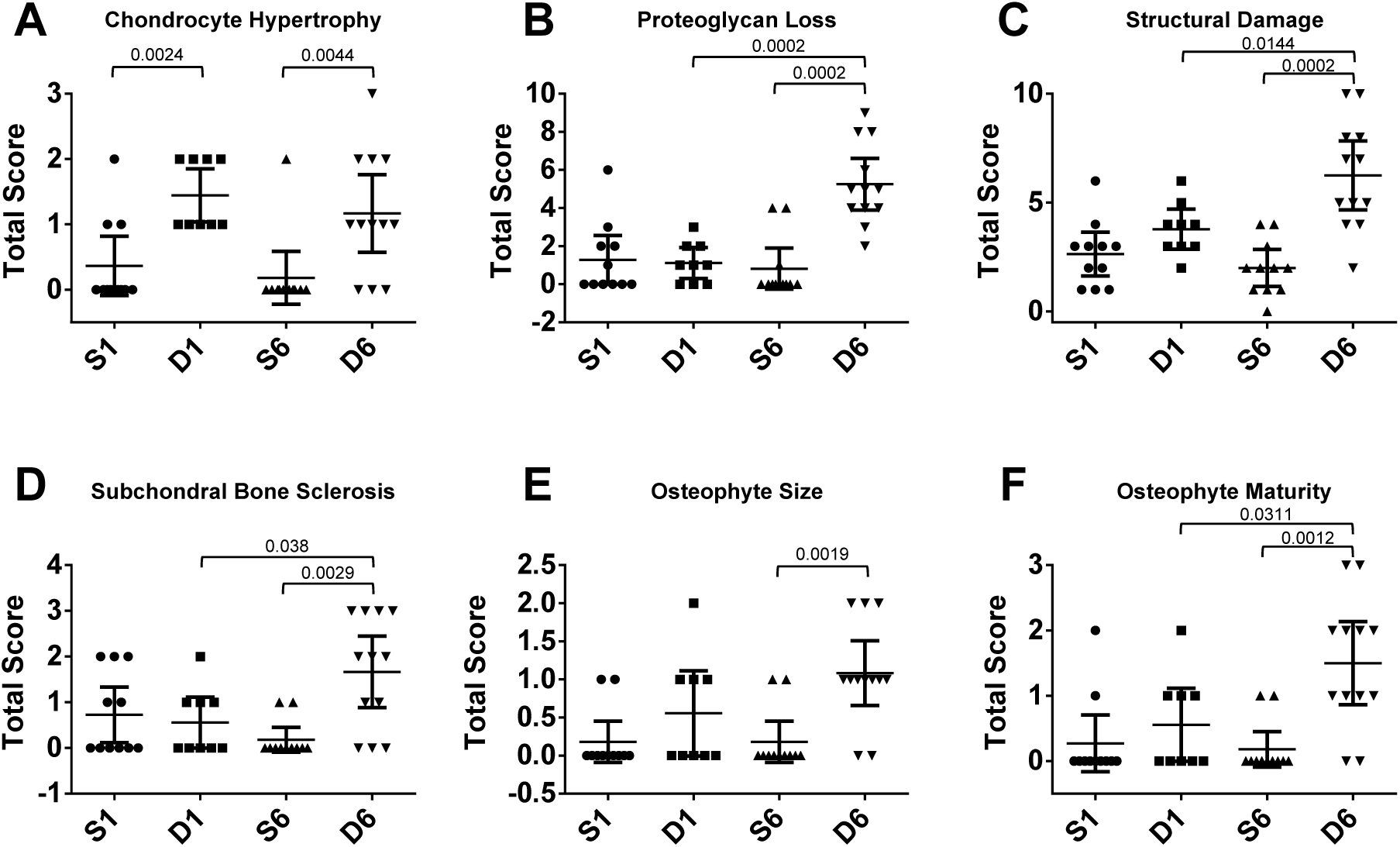
Cumulative histopathological scores of **(A)** chondrocyte hypertrophy, **(B)** cartilage proteoglycan loss, **(C)** articular cartilage structural damage, **(D)** subchondral bone sclerosis and osteophyte **(E)** size and **(F)** maturity in the proximal tibia of wild type sham and DMM operated knee joints at 1 and 6 weeks post surgery. Scatter plots display values for each individual mouse and mean (horizontal bar) ± 95% confidence intervals. Significant differences between groups connected by lines with exact p-values for each comparison indicated above the line, as determined by Mann-Whitney U-test with Benjamini-Hochberg multiple comparison correction. n = 11 (S1), n = 9 (D1), n = 11 (S6), n = 12 (D6). S = Sham; D = DMM.

There was no difference in SCB sclerosis between surgeries at 1 week (Fig. 1D) but by week 6 there was significant sclerosis in DMM compared with sham (p = 0.003) and in DMM at 6 weeks compared with DMM at 1 week (p = 0.038). There was no osteophyte development at 1 week but at 6 weeks they had formed in DMM only, being larger (p = 0.002) and more mature (p = 0.001) than sham (Fig. 1E&F). Osteophyte maturation from cartilage to bone also progressed with time in DMM joints (p=0.03).

### miR expression profiling of osteoarthritic SCB tissue

To assess the role of miRs in the early and progressive stages of OA we performed miR microarray analyses on 1 and 6 week post sham and DMM SCB and cartilage samples. Of the 1,881 miRs represented on the Agilent microarrays, 490 miRs were detected above background in SCB samples. Following stringent statistical analysis as described previously (Kung et al., 2017), we did not observe any statistically significant differences in miR expression in SCB between DMM and sham surgeries at either 1 or 6 week time points (Fig. 2A,B; data available online at www/ncbi.nlm.nih.gov/geo/; accession number GSE93008). However, there were statistically significant miR expression changes with post-operative time in SCB tissues (Supplemental Table 1). 37 miRs were dysregulated between 1 and 6 week time points in SCB (adj.p.value < 0.05; Supplemental Table 1) demonstrating the sensitivity and confidence of the microarray approach to detect miR expression changes if present.

**Figure 2.**
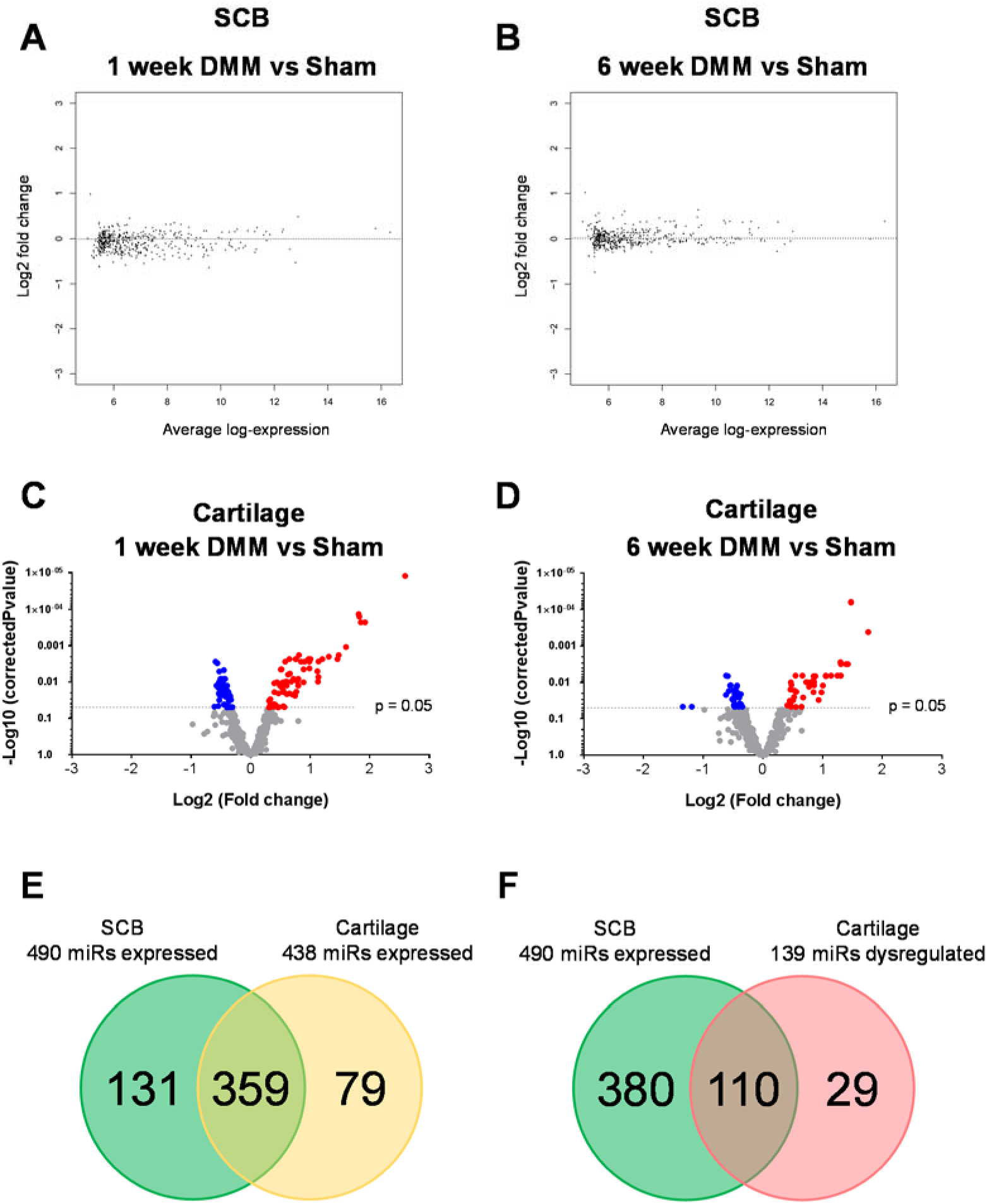
SCB and articular cartilage miR expression profiling. MA plots displaying average log-expression intensity values and log2 fold change for each miR expressed in SCB between DMM and sham in (A) 1 week and (B) 6 weeks. There were no significant changes in miR expression in SCB of DMM vs sham joints. Volcano plots showing differentially expressed miRs with adj.p-value < 0.05 in cartilage at (C) 1 and (D) 6 weeks post DMM vs sham. (C) 122 miRs were significantly dysregulated (adj.p-value < 0.05) in 1 week DMM vs sham (highlighted in blue+red). Of those, 69 miRs were up-regulated with adj.p-value < 0.05 (highlighted in red) and 53 miRs were down-regulated with adj.p-value < 0.05 (highlighted in blue) in 1 week DMM vs sham. (D) 74 miRs were significantly dysregulated (adj.p-value < 0.05) in 6 week DMM vs sham adj.p-value < 0.05 (highlighted in blue+red). Of those, 42 miRs were up-regulated with adj.p-value < 0.05 (highlighted in red) and 32 miRs were down-regulated with adj.p-value < 0.05 (highlighted in blue) in 6 week DMM vs sham. (E) Comparison of miR expression in SCB and articular cartilage. (F) 110 out of 139 miRs dysregulated in cartilage were also expressed but not dysregulated in SCB. 29 miRs were uniquely expressed and dysregulated in cartilage.

### Dysregulated miR expression in DMM-induced OA cartilage

438 miRs were detected above background in cartilage samples. Of these, 359 miRs (~80%) were also expressed in SCB, with the remaining 79 miRs exclusively expressed in cartilage (Fig. 2E). In contrast to the SCB above, miR expression profiling revealed 122 and 74 miRs that were significantly differentially expressed (adj.p.value < 0.05) in cartilage of DMM versus sham-operated mice at 1 and 6 weeks post-surgery, respectively (Supplemental Tables 2-3). Of those, 69 miRs were up-regulated and 53 miRs were down-regulated in 1 week DMM versus sham (Fig. 2C). At 6 weeks, 42 miRs were up-regulated and 32 miRs were down-regulated in DMM versus sham (Fig. 2D).

Further analysis also demonstrated temporal patterns of differential miR expression. Of the 139 miRs dysregulated in total during the 6 week study, 65 and 17 miRs were uniquely dysregulated at 1 and 6 weeks, respectively (Fig. 3A(i),B), signifying these as potentially involved in OA initiation (week 1) and the development of cartilage degeneration over the first 6 weeks. Additionally, a cohort of 57 dysregulated miRs were common to both 1 and 6 week time points in the array data (Fig. 3B). Amongst this 57, 12 miRs were identified with a robust array fold change > 2.0 and adj.p.value < 0.05, including miR-6931-5p, miR-466i-5p, miR-3082-5p, miR-1187, miR-669n, miR-468-3p, miR-669l-5p, miR-669e-5p and miR-672-5p, miR-574-5p, miR-32-3p (Supplemental Tables 2-3). Moreover, of the 139 dysregulated in cartilage in total, 110 miRs were found to be also co-expressed in SCB but not differentially regulated. The remaining 29 dysregulated miRs were found to be unique to cartilage (Fig. 2F). This segregation of differential expression indicates the presence of tissue specific dysregulation of miRs, the majority of which were commonly expressed between the two tissues.

**Figure 3.**
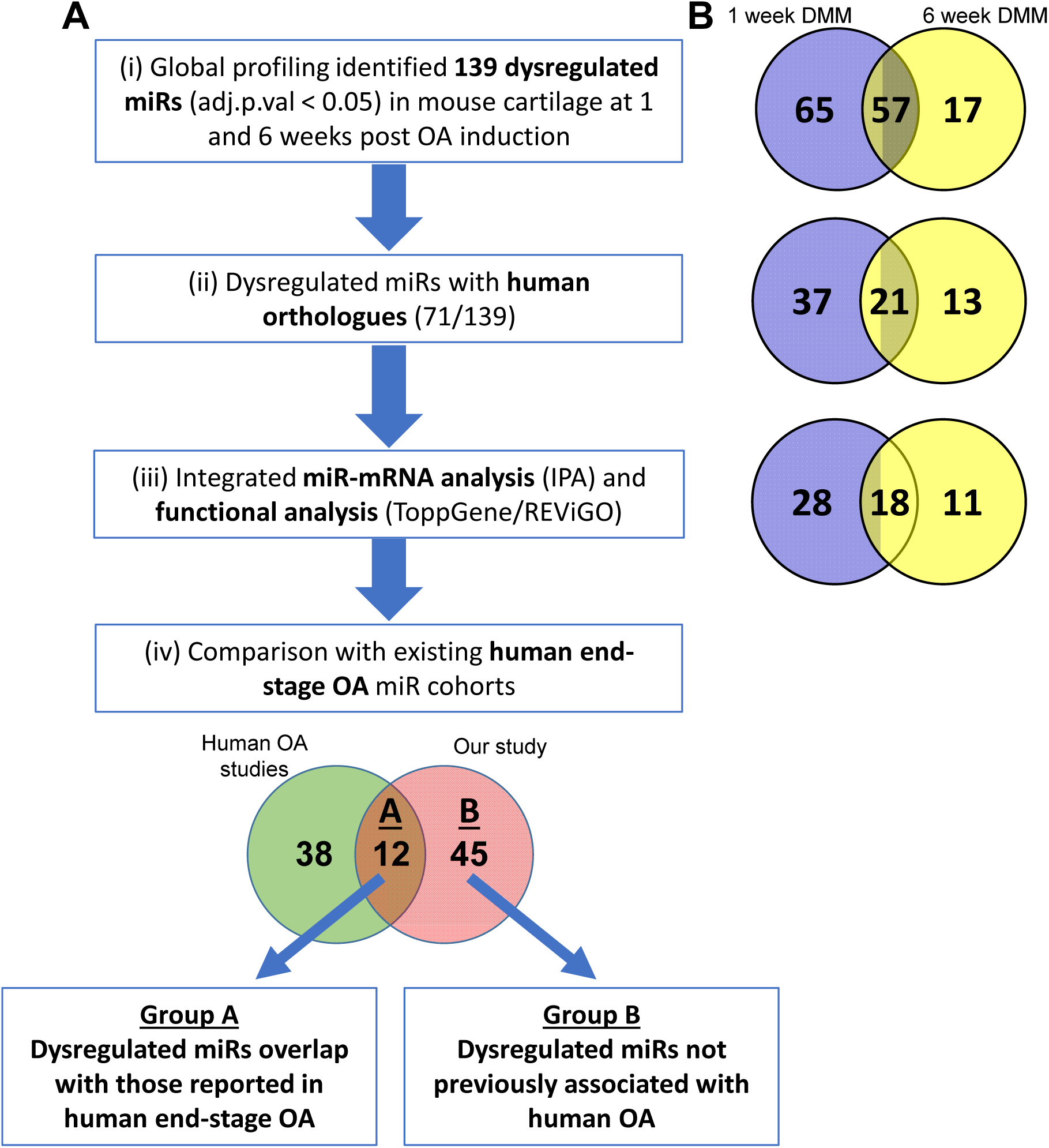
Comprehensive unbiased candidate miR filtering strategy. **(A)** Flow chart of the filtering criteria employed to select candidate miRs. **(B)** Venn diagrams comparing dysregulated miRs at 1 and 6 week DMM following each filtered criteria. **(i)** Statistically significant miRs dysregulated in either 1 and/or 6 weeks DMM compared with sham (139 miRs in total) were filtered based on **(ii)** the existence of their known human orthologues (71/139). **(iii)** mRNA (target) expression data was integrated alongside miR expression data in Ingenuity based target prediction analysis (IPA). 57 miRs displayed target mRNA information which was subsequently used in downstream functional analysis. **(iv)** Lastly, these 57 miRs were compared with miR candidates previously identified in human end-stage OA studies. 12 miRs overlapped with those already described in human end-stage OA (Group A) and 45 miRs include novel miR candidates not previously identified in human OA (Group B).

### Comprehensive filtering strategy to select candidate miRs

Successful functional analysis and therapeutic testing of dysregulated miRs in future studies relies critically on the selection of suitable miR candidates. Therefore, to select miR candidates from the pool of 139 dysregulated miRs (either at 1 and/or 6 weeks) we employed a stringent and comprehensive filtering approach outlined in Figure 3. Firstly, dysregulated miRs were filtered (based on name and/or mature sequence homologies) to select those with known human orthologues. This defined 71 out of 139 dysregulated miRs with existing human miR orthologues (Fig. 3A(ii)), thus more likely to be of human disease relevance. Secondly, to gain an insight in the biological function of these miRs, mRNA expression data from the same model and time points (Bateman et al., 2013) were integrated with the miR expression data to identify miRs that potentially impact OA pathological gene networks (Fig. 3A(iii); see below). miRs with no mRNA target data were excluded from further analysis. Lastly, our filtered dataset was compared with findings from four independent human end-stage OA miR studies (Diaz-Prado et al., 2012; Iliopoulos et al., 2008; Jones et al., 2009; Song et al., 2013). From this, two cohorts of miR candidates were generated (Fig. 3A(iv)); Group A - Dysregulated miRs that overlap with human end-stage OA miR candidates and Group B - miRs including novel dysregulated miRs not previously described in OA (see below).

### Identification of putative target genes by integrated miR:mRNA expression analysis

To drill down on the functional importance of OA-dysregulated miRs that have known human orthologues we performed a bioinformatics-based analysis to identify putative target genes significantly regulated by miRs differentially expressed in OA cartilage at 1 and 6 weeks post DMM (Supplemental Tables 4-5). We integrated both miR and mRNA datasets by using the microRNA target prediction module within Ingenuity Pathway Analysis (IPA) which utilises computational algorithms and databases (including TargetScan Human, Tarbase) to predict or display experimentally observed target genes. To do this we utilised our previous global mRNA expression profiling of cartilage from DMM-operated WT mice (at the same 1 and 6 week time points) (Bateman et al., 2013). The analysis recognised and correlated the expression of miRs with mRNA expression (targets) and filtered inversely dysregulated miR-mRNA interactions, many of which had high relevance to OA pathology (Supplemental Tables 4-5). At 1 week DMM, IPA miR target analysis recognised 46 miRs targeting 397 mRNAs and at 6 weeks there were 29 miRs targeting 510 mRNAs (Supplemental Tables 4-5). Altogether, mRNA target data was generated for 57 of the 71 differentially expressed miRs with known human orthologues across both time points (Fig. 3A(iii)).

For example, *Ptgs2* (*Cox-2*) which was one of the most highly up-regulated genes in DMM cartilage at both 1 and 6 weeks (Bateman et al., 2013), has been shown to be experimentally regulated by multiple miRs in our dataset, including miR-98-5p, miR-26b-5p, miR-15a-5p/miR-16-5p and Let-7d-5p, all of which were statistically significantly down-regulated in DMM cartilage (Supplemental Tables 4-5; Table 1&2). Similarly, *Phlda2* the most up-regulated gene at 1 week, may be a putative target of miR-128-3p, miR-22-5p and miR-376-3p (Supplemental Tables 4-5).

**Table 1.** IPA miR target filter analysis on integrated miR:mRNA expression data in 1 week DMM. Predicted and/or experimentally validated mRNA targets of Group A dysregulated miRs that overlap with human end-stage OA miR candidates.

| miR | Fold Change | Source | Confidence | Target Gene | Fold Change |
| --- | --- | --- | --- | --- | --- |
| miR-26b-5p | -1.27 | Ingenuity Expert Findings, TargetScan Human | Experimentally Observed, Moderate (predicted) | PTGS2 | 19.16 |
|  |  | TargetScan Human | Moderate (predicted) | COL10A1 | 9.68 |
|  |  | TargetScan Human | High (predicted) | HMGA1 | 4.019 |
|  |  | TargetScan Human | Moderate (predicted) | SAR1B | 3.415 |
|  |  | TargetScan Human | High (predicted) | SLC38A2 | 3.065 |
| miR-30c-5p | -1.46 | TarBase, TargetScan Human | Experimentally Observed, Moderate (predicted) | NT5E | 14.162 |
|  |  | TargetScan Human | Moderate (predicted) | ACSBG1 | 8.129 |
|  |  | TarBase | Experimentally Observed | NPR3 | 7.306 |
|  |  | TargetScan Human | Moderate (predicted) | TUSC3 | 4.411 |
|  |  | TargetScan Human | High (predicted) | MEX3B | 4.383 |
| miR-98-5p | -1.41 | TarBase | Experimentally Observed | PTGS2 | 19.16 |
|  |  | TargetScan Human | Moderate (predicted) | CHSY3 | 6.22 |
|  |  | TargetScan Human | High (predicted) | NGF | 4.807 |
|  |  | TargetScan Human | Moderate (predicted) | HBEGF | 4.754 |
|  |  | Ingenuity Expert Findings | Experimentally Observed | HMGA2 | 4.405 |
| miR-149-5p | -1.24 | TargetScan Human | Moderate (predicted) | NPR3 | 7.306 |
|  |  | TargetScan Human | Moderate (predicted) | TNFRSF12A | 5.946 |
|  |  | TargetScan Human | Moderate (predicted) | DLL1 | 5.359 |
|  |  | TargetScan Human | Moderate (predicted) | FOSL1 | 4.708 |
|  |  | TargetScan Human | Moderate (predicted) | KLHL21 | 4.374 |
| miR-210-3p | -1.26 | TargetScan Human | High (predicted) | MEX3B | 4.383 |
|  |  | TargetScan Human | Moderate (predicted) | EPB41L1 | 3.202 |
|  |  | TargetScan Human | Moderate (predicted) | TRIB1 | 2.711 |
| miR-337-3p | -1.31 | TargetScan Human | Moderate (predicted) | FOS | 10.091 |
|  |  | TargetScan Human | Moderate (predicted) | FGF1 | 5.404 |
|  |  | TargetScan Human | Moderate (predicted) | RGS16 | 4.031 |
|  |  | TargetScan Human | Moderate (predicted) | CD44 | 4.008 |
|  |  | TargetScan Human | Moderate (predicted) | ZZZ3 | 3.706 |

**Table 2.**
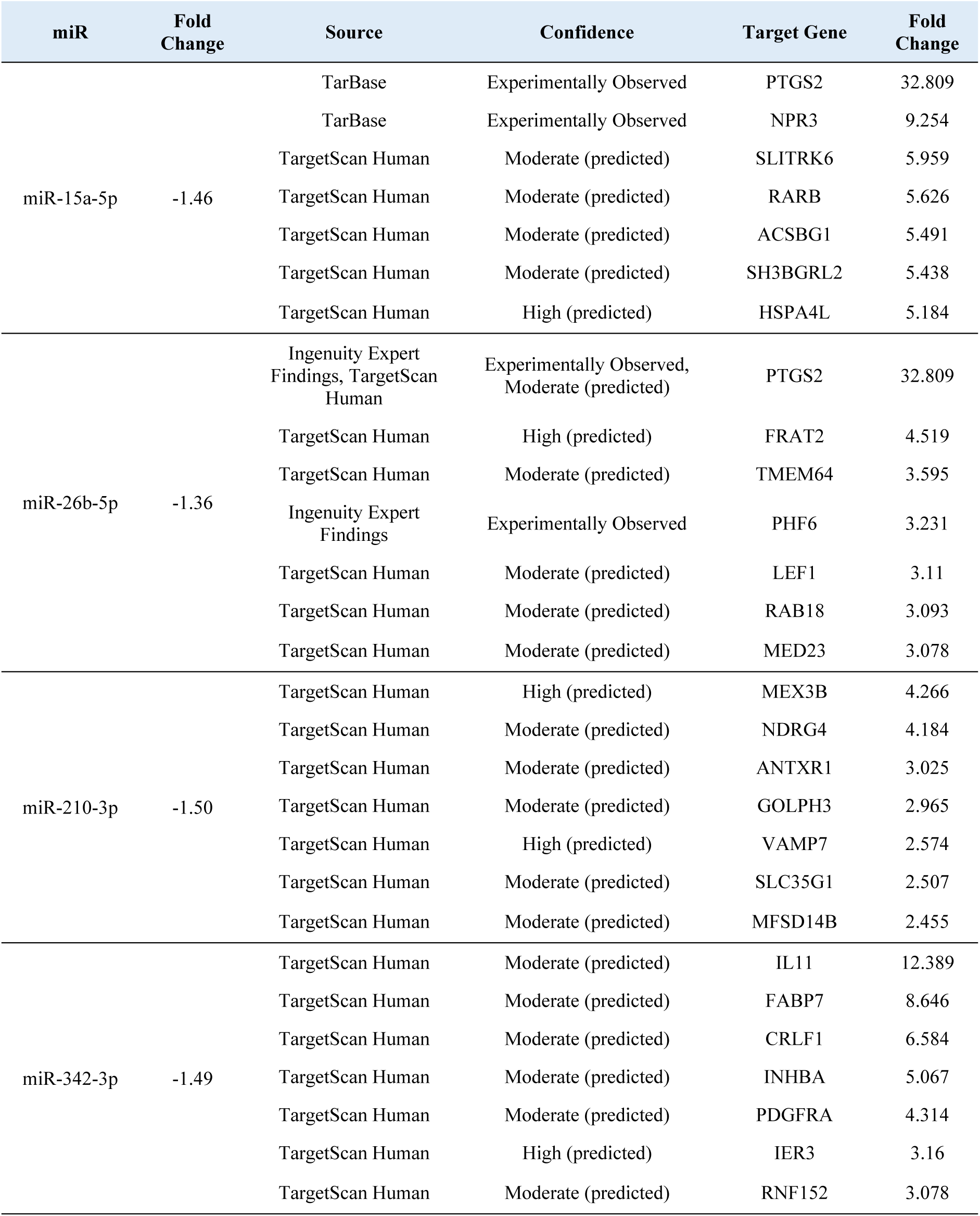
IPA miR target filter analysis on integrated miR:mRNA expression data in 6 week DMM. Predicted and/or experimentally validated mRNA targets of Group A dysregulated miRs that overlap with human end-stage OA miR candidates.

| miR | Fold Change | Source | Confidence | Target Gene | Fold Change |
| --- | --- | --- | --- | --- | --- |
| miR-15a-5p | -1.46 | TarBase | Experimentally Observed | PTGS2 | 32.809 |
|  |  | TarBase | Experimentally Observed | NPR3 | 9.254 |
|  |  | TargetScan Human | Moderate (predicted) | SLITRK6 | 5.959 |
|  |  | TargetScan Human | Moderate (predicted) | RARB | 5.626 |
|  |  | TargetScan Human | Moderate (predicted) | ACSBG1 | 5.491 |
|  |  | TargetScan Human | Moderate (predicted) | SH3BGRL2 | 5.438 |
|  |  | TargetScan Human | High (predicted) | HSPA4L | 5.184 |
| miR-26b-5p | -1.36 | Ingenuity Expert Findings, TargetScan Human | Experimentally Observed, Moderate (predicted) | PTGS2 | 32.809 |
|  |  | TargetScan Human | High (predicted) | FRAT2 | 4.519 |
|  |  | TargetScan Human | Moderate (predicted) | TMEM64 | 3.595 |
|  |  | Ingenuity Expert Findings | Experimentally Observed | PHF6 | 3.231 |
|  |  | TargetScan Human | Moderate (predicted) | LEF1 | 3.11 |
|  |  | TargetScan Human | Moderate (predicted) | RAB18 | 3.093 |
|  |  | TargetScan Human | Moderate (predicted) | MED23 | 3.078 |
| miR-210-3p | -1.50 | TargetScan Human | High (predicted) | MEX3B | 4.266 |
|  |  | TargetScan Human | Moderate (predicted) | NDRG4 | 4.184 |
|  |  | TargetScan Human | Moderate (predicted) | ANTXR1 | 3.025 |
|  |  | TargetScan Human | Moderate (predicted) | GOLPH3 | 2.965 |
|  |  | TargetScan Human | High (predicted) | VAMP7 | 2.574 |
|  |  | TargetScan Human | Moderate (predicted) | SLC35G1 | 2.507 |
|  |  | TargetScan Human | Moderate (predicted) | MFSD14B | 2.455 |
| miR-342-3p | -1.49 | TargetScan Human | Moderate (predicted) | IL11 | 12.389 |
|  |  | TargetScan Human | Moderate (predicted) | FABP7 | 8.646 |
|  |  | TargetScan Human | Moderate (predicted) | CRLF1 | 6.584 |
|  |  | TargetScan Human | Moderate (predicted) | INHBA | 5.067 |
|  |  | TargetScan Human | Moderate (predicted) | PDGFRA | 4.314 |
|  |  | TargetScan Human | High (predicted) | IER3 | 3.16 |
|  |  | TargetScan Human | Moderate (predicted) | RNF152 | 3.078 |

In addition, we identified other potentially important miR-mRNA paired interactions involved in OA pathological networks, such as, miR-26b-5p-*Col10a1*, miR-224-5p/miR-425-5p-*Il*-*11*, miR-28-5p/miR-30c-5p/miR-377-3p-*Nt5e*, miR-337-3p-*Fos*, miR-128-3p-*Npy* and *Npr3* potentially regulated by several miRs, including miR-107-3p, miR-10a-5p, miR-128-3p, miR-15a-5p, miR-30c-5p, miR-196b-5p and miR-411-3p (Supplemental Tables 4-5; Table 1&2).

### Comparison of our dataset with previously published human OA cartilage studies

Several dysregulated miRs have been identified in human end-stage OA-cartilage or chondrocytes from previous expression profiling experiments (Diaz-Prado et al., 2012; Iliopoulos et al., 2008; Jones et al., 2009; Song et al., 2013). Differentially expressed miRs identified from these 4 independent studies were compared, with related miR family members allocated into the same group as they most likely target the same genes/pathways e.g. miR-15a-5p and miR-16-5p, miR-26a/b, miR-30b/c/d. Nonetheless, there was a striking lack of overlap with only 6 out of 50 miRs identified (miR-342-3p, miR-25, miR-22-3p, miR-23b, miR-26-5p, miR-30-5p) common between any of the human studies (Fig. 4).

**Figure 4.**
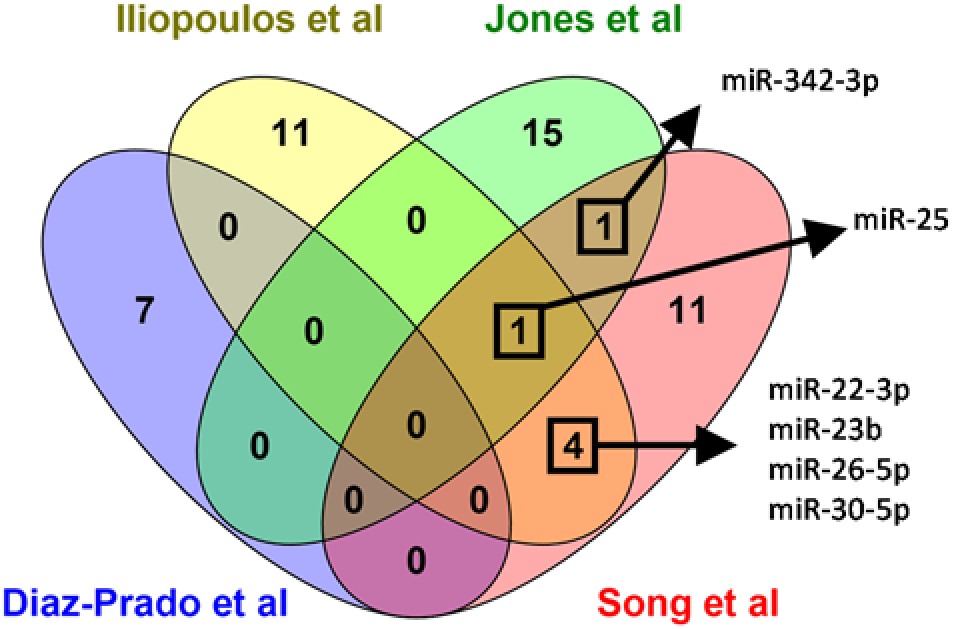
miR expression meta-analysis comparing the amount of overlap between previously published human end-stage OA studies. Differentially expressed miRs identified from 4 independent studies examining end-stage human OA cartilage/chondrocytes were compared against one another. Only 6 out of 50 dysregulated miRs were common between more than one study.

In contrast, of the 50 dysregulated human OA miRs (Diaz-Prado et al., 2012; Iliopoulos et al., 2008; Jones et al., 2009; Song et al., 2013), 12 miRs (miR-15a/16-5p, miR-26b-5p, miR-30c-5p, miR-98-5p, miR-107-3p, miR-149-5p, miR-185-5p, miR-210-3p, miR-223-3p, miR-337-3p, miR-342-3p and miR-377-3p) overlapped with our current study (Group A; Fig. 3A(iv)). Furthermore, all 12 of these Group A miR candidates displayed OA-regulated mRNA target data (Supplemental Tables 4-5; Table1&2). IPA-generated target filter analysis revealed several interesting miR-mRNA interactions for this candidate group. For instance, miR-26b-5p-*Ptgs2/-Col10a1*, miR-30c-5p-*Nt5e/-Npr3*, miR-98-5p-*Ptgs2/-Chsy3/-Ngf*, miR-149-5p-*Npr3/-Tnfrsf12a/-Dll1/-Fosl*, miR-337-3p-*Fos/-Fgf1*, miR-15a-5p-*Ptgs2/-Npr3* and miR-342-3p-*Il-11/-Crlf1/-Inhba/-Pdgfra* (Tables 1&2).

The remaining 45 miRs (Group B; Fig. 3A(iv)) in our filtered dataset didn’t overlap with the human studies and have not been previously associated with OA (based on these 4 studies). Of these Group B miRs which include novel miR candidates, the most up-regulated were miR-574-5p, miR-31-5p and miR-195a-3p and the most down-regulated were miR-196b-5p, miR-411-3p and miR-451a (Supplemental Tables 2-3; Group B miRs highlighted in yellow). Within this Group B, Let-7d-5p was an interesting candidate with *Ptgs2* as an experimentally observed target (Supplemental Table 5). IPA-generated target filter analysis also revealed *Tns1* (Tensin-1), a focal adhesion protein, as a potential target of miR-31-5p (Supplemental Tables 2-3).

### qPCR validation of miR candidates

miRs candidates in Group A and B prioritized through our filtering strategy were tested by qPCR to confirm their dysregulation in OA on additional biological replicates (Fig. 5&6). Of the miRs in Group A, 7 miRs were confirmed as differentially regulated in wild type DMM compared with sham at 1 and/or 6 week time points (miR-15/-16-5p, miR-26b-5p, miR-30c-5p, miR-98-5p, miR-149-5p, miR-210-3p and miR-342-3p; Fig. 5). No statistically significant differences were observed for miR-107-3p, miR-185-5p and miR-223-3p in this larger cohort of mice (Fig. 5). miR-337-3p and miR-377-3p were below the level of detection by qPCR.

**Figure 5.**
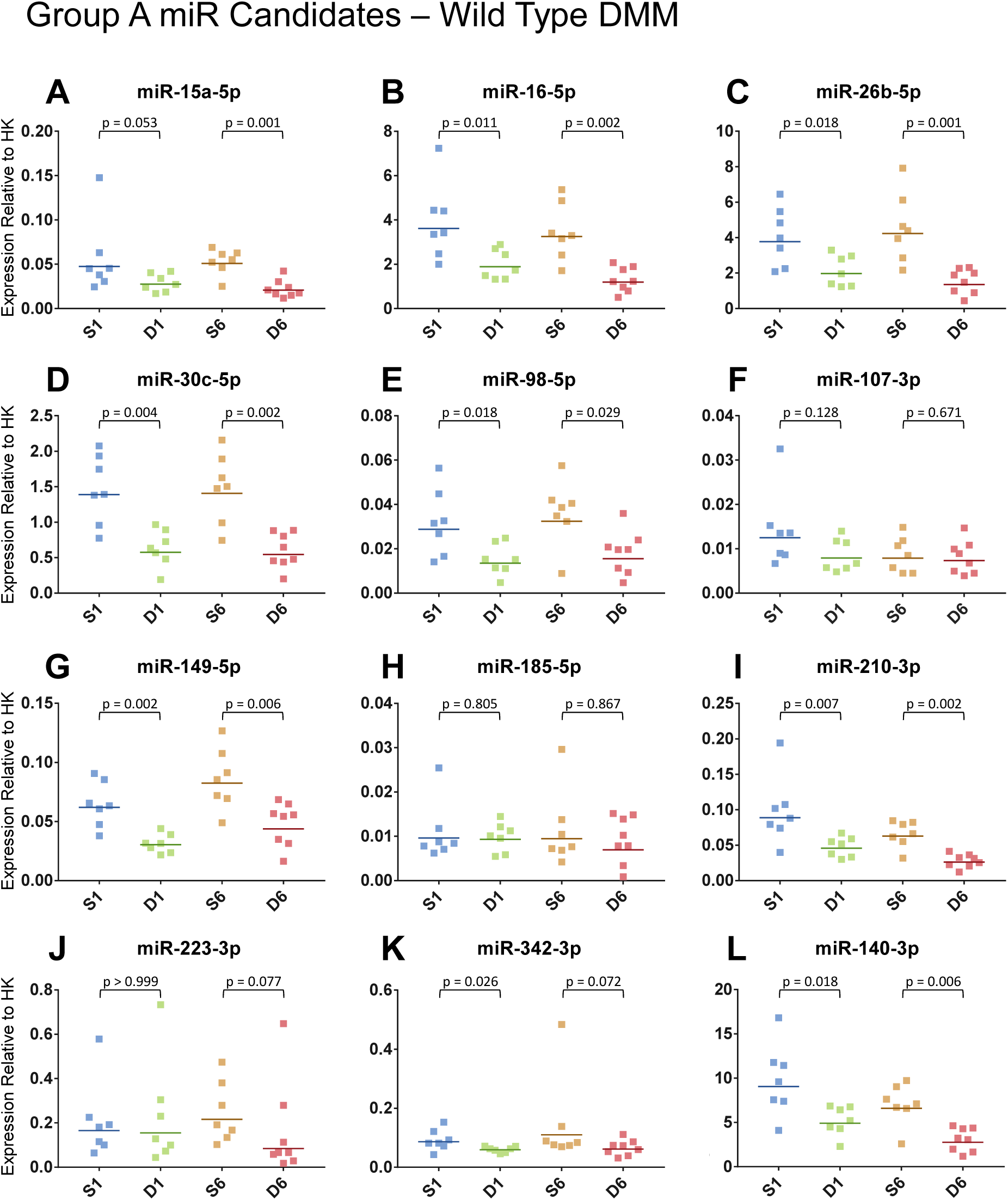
Validation of Group A miR candidates by qPCR. Expression of miRs in articular cartilage tissue from wild type mice at 1 and 6 weeks post sham or DMM surgery was determined. Data shown as expression relative to the average expression of two housekeepers (HK). Each symbol represents an individual mouse (average of 2 technical replicates). Horizontal bars show the geometric mean expression (n=7-8/surgery/timepoint). Statistical differences as determined by Mann-Whitney U test between groups connected by lines with exact p-values for each comparison indicated above the line. S = Sham; D = DMM.

A subset of novel miRs candidates in Group B were also examined by qPCR to validate their dysregulation in OA DMM cartilage (Fig. 6). miR-574-5p and miR-31-5p were confirmed as significantly up-regulated in wild type DMM cartilage and Let-7d-5p was significantly down-regulated in wild type DMM cartilage (Fig. 6A-C), thus confirming the microarray data. No statistically significant differences was observed for miR-451a in the larger cohort of mice (Fig. 6D) and miR-195a-3p and miR-411-3p were undetectable by qPCR.

**Figure 6.**
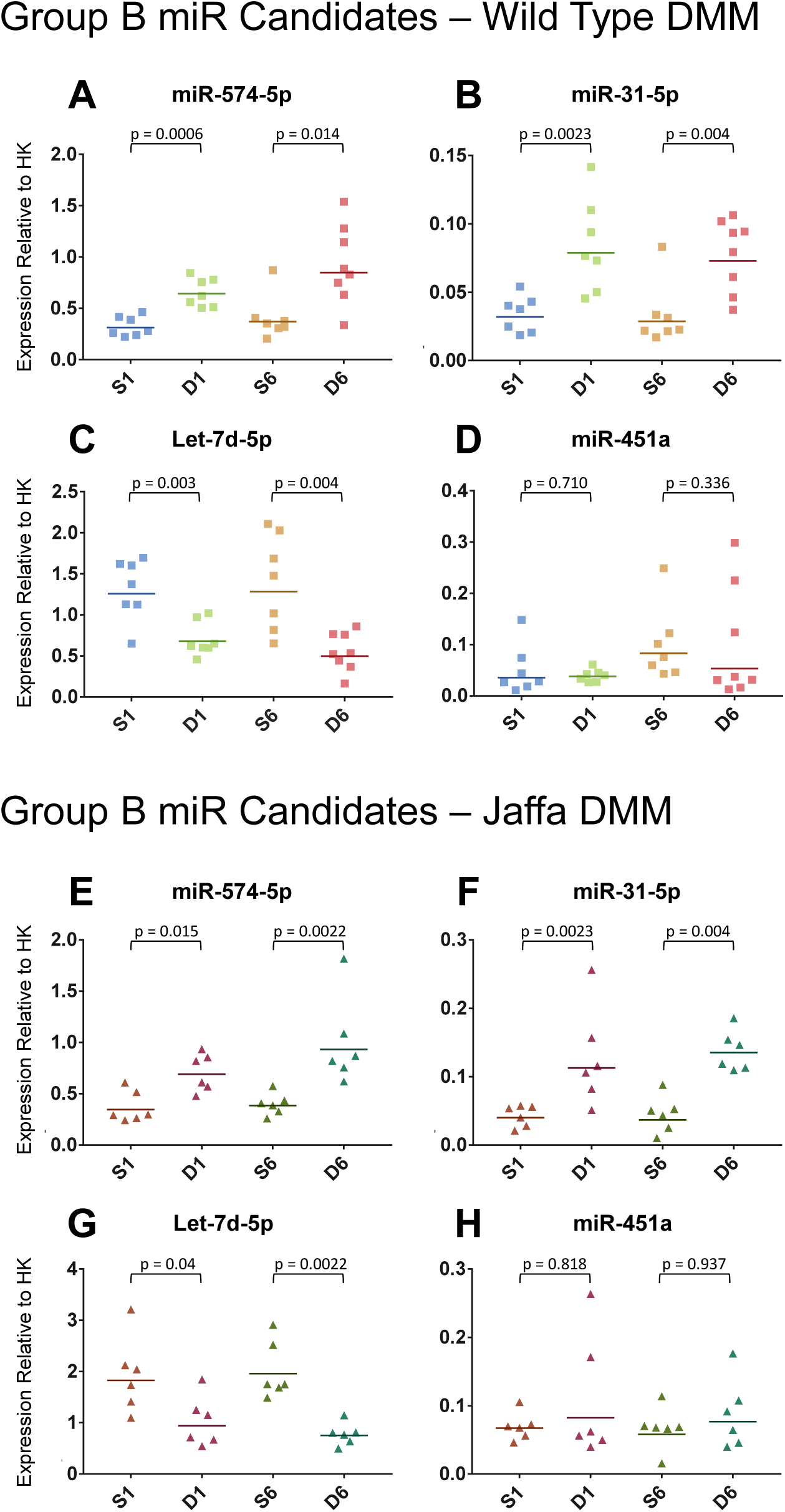
Validation of Group B miR candidates by qPCR. Expression of miRs in articular cartilage tissue from wild type mice (A-D) and in mice resistant to aggrecan breakdown (Jaffa; E-H) at 1 and 6 weeks post sham or DMM surgery was determined. Data shown as expression relative to the average expression of two housekeepers (HK). Each symbol represents an individual mouse (average of 2 technical replicates). Horizontal bars show the geometric mean expression (n=6-8/surgery/timepoint). Statistical differences as determined by Mann-Whitney U test between groups connected by lines with exact p-values for each comparison indicated above the line. S = Sham; D = DMM.

We also examined the expression of miR-140 in this larger cohort as it is known to be an important cartilage-specific miR involved in maintaining cartilage homeostasis (Miyaki et al., 2010). qPCR analysis revealed a significant decrease in miR-140 expression in DMM-operated cartilage at both 1 and 6 weeks in comparison to sham controls (Fig. 5), thus corroborating with previous findings in human OA cartilage (Iliopoulos et al., 2008; Miyaki et al., 2009; Song et al., 2013) and further demonstrating the validity of the DMM surgery as a model of human OA.

### Dysregulated miR expression independent of aggrecan breakdown

Finally, we interrogated the expression of our validated Group A and B miR candidates in mice carrying a knockin mutation in aggrecan which endows resistance to aggrecanase cleavage in the interglobular domain (IGD) and protection from DMM-induced cartilage proteoglycan loss and subsequent structural damage (Jaffa; *Acan* p.374ALGS→374NVYS) (Little et al., 2007). Identifying miRs dysregulated in both wild type and Jaffa DMM cartilage allowed us to discriminate between miRs which are independent of aggrecan breakdown with those that are not dysregulated in Jaffa DMM and thus dependent on the degradation of aggrecan. Interestingly, dysregulated candidate miRs were on-the-whole similarly dysregulated in Jaffa DMM mice (Fig. 6&7). miR-15a-5p, miR-16-5p, miR-26b-5p, miR-30c-5p, miR-98-5p, miR-210-3p, miR-342-3p, miR-140 and Let-7d-5p were statistically significantly down-regulated in Jaffa DMM (Fig. 6&7) mice in a similar fashion as wild type DMM mice (Fig. 5&6). miR-149-5p was down regulated in Jaffa at 1 week but not 6 weeks (Fig. 7F). miR-574-5p and miR-31-5p were also significantly up-regulated in Jaffa DMM mice in a similar fashion as wild type DMM mice (Fig. 6E,F).

**Figure 7.**
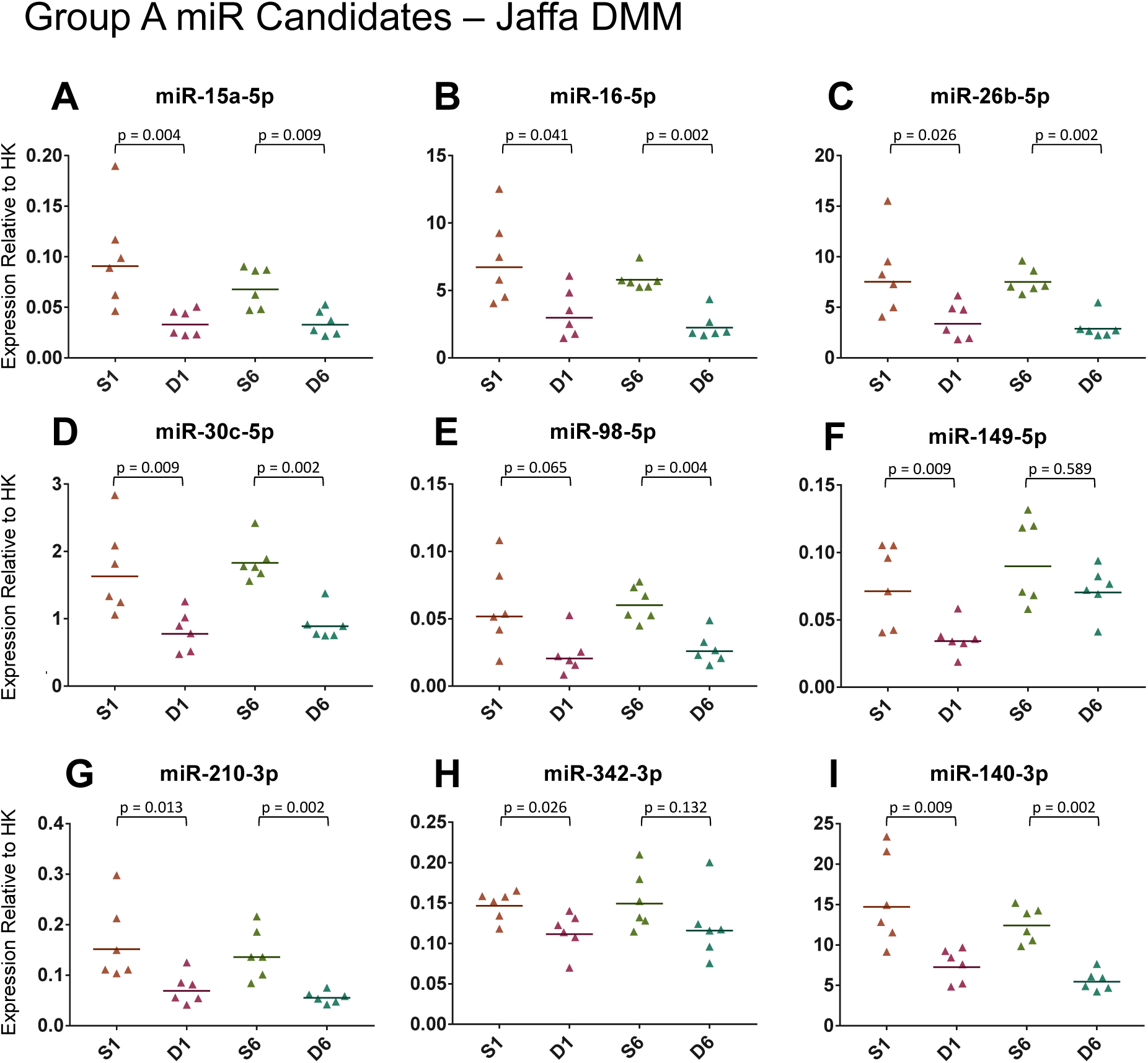
qPCR analysis of Group A miR candidates in mice resistant to aggrecan breakdown (Jaffa). Expression of miRs in cartilage from Jaffa mice at 1 and 6 weeks post sham or DMM surgery was determined. Data shown as expression relative to the average expression of two housekeepers (HK). Each symbol represents an individual mouse (average of 2 technical replicates). Horizontal bars show the geometric mean expression (n=6/surgery/timepoint). Statistical differences as determined by Mann-Whitney U test between groups connected by lines with exact p-values for each comparison indicated above the line. S = Sham; D = DMM.

## DISCUSSION

In this study, we have profiled miR expression in cartilage and SCB tissue of early OA joints induced by DMM surgery with the aim of exploring novel OA disease mechanisms which could underpin new therapeutic strategies. DMM mice demonstrated both time-dependent (cartilage proteoglycan loss, structural damage, SCB sclerosis, osteophytes) and independent (chondrocyte hypertrophy) OA pathology, replicating key pathological features of the human condition. Correspondingly, we observed changes in miR expression specifically in cartilage that mirrored the temporal changes of proteoglycan loss and structural damage (specific to 6 weeks DMM) and the static changes of chondrocyte hypertrophy (both 1 and 6 weeks DMM). Consistent with the early detection of chondrocyte hypertrophy, *Col10a1* was previously shown to be significantly up-regulated in 1 week DMM (Bateman et al., 2013).

We found that in total 139 miRs were dysregulated in cartilage during the development of post-traumatic OA over 6 weeks. Interestingly, 122 of these dysregulated miRs (~88%) were identified prior to any significant aggrecan degradation or cartilage structural degeneration at 1 week DMM suggesting that significant miR dysregulation could be involved in the initiation of OA pathology. Moreover, miR dysregulation prior to pathology represents an ideal window for diagnostic and therapeutic benefit for patients following post-traumatic injury. Importantly, we interrogated the expression of DMM-dysregulated candidate miRs in mice resistant to aggrecan cleavage in the IGD which prevents the release of the entire glycosaminoglycan-containing portion of aggrecan by aggrecanases. This allowed us to identify miR expression changes that are dependent on, versus those that are independent of, ADAMTS-driven aggrecanolysis. All of the candidate miRs examined were shown to be independent of aggrecan cleavage, consistent with the dysregulation of these miRs prior to signs of proteoglycan loss and structural damage in wild type DMM-operated mice. This suggests that the differential expression of these miRs occurs not as a result of the inflammation induced by joint injury/surgery (equivalent in sham and DMM) but as a response to increased mechanical stress induced by DMM surgery and the acute-initiation of the OA process rather than the downstream consequences of aggrecan breakdown. While robust functional analysis of the targets of the dysregulated candidate miRs will be needed to tease out key miRs that definitively relate to OA pathology, these miR candidates can be segregated into several cohorts to explore possible roles in the initiation and progression of cartilage damage in human OA.

Firstly, to maximize the translational relevance to patients, all OA-dysregulated cartilage miRs were filtered to select those with human orthologues. This defined 71 out of 139 miRs dysregulated in total. The relevance of the dysregulated miRs without human orthologues (68/139) is unclear. Currently, there are 2588 mature miRs known in humans but new ones are continuously being discovered. New evidence has emerged suggesting that over 1000 novel human miRs are yet to be uncovered as sequencing of more tissues and advancing technology come into play (Friedländer et al., 2014). Accordingly, deep sequencing was performed to identify novel miRs in human OA cartilage (Crowe et al., 2016). Novel candidate miRs were characterized and validated in human tissue panels, chondrogenesis, chondrocyte differentiation and cartilage injury models, which led to the identification of 3 novel miRs with possible functional roles in cartilage homeostasis and OA (Crowe et al., 2016). Current studies suggest that we are only scraping the surface of miR biology and it is likely that more are to be discovered and annotated. Of the group of miRs lacking human counterparts but were significantly up-regulated in DMM cartilage at both time points (e.g. miR-6931-5p, miR-466i-5p, miR-3082-5p, miR-1187), most have not been associated with OA previously and their roles are not well known. miR-466i-5p on the other hand has been suggested to have a role in controlling inflammation, through targeting the expression of inflammatory markers such as Cox-2/Ptgs2 and iNOS in liver inflammation (Saravanan et al., 2015). miR-466i-5p has also been implicated in hedgehog signalling, PDGF signalling, response to stress pathways and Wnt pathway in the context of renal disease (Cheng et al., 2013). The roles of these miRs specifically in OA and their relevance to the human condition remains to be determined.

To gain an insight into the biological function of the remaining miRs with human counterparts and thus identify high-priority miR candidates for future study, we conducted miR target analysis (IPA) by integrating the miR expression data with our previous mRNA expression data collected from the same DMM model and time points. This allowed us to focus our study on miRs that are most likely to target OA-pathological networks and pathways. This further shortlisted our cohort to 57 miRs (Fig. 3Aiii). In addition, we compared our dataset with current miR datasets from human OA (Diaz-Prado et al., 2012; Iliopoulos et al., 2008; Jones et al., 2009; Song et al., 2013). Notwithstanding, that our OA model coincides with early stage cartilage degeneration and the human samples are from end-stage disease cartilage, miRs common to both groups are thus likely to be important throughout the OA process. We identified and confirmed a group of 7 dysregulated miRs candidates (termed Group A; miR-15a/16-5p, miR-26b-5p, miR-30c-5p, miR-98-5p, miR-149-5p, miR-210-3p, miR-342-3p) of particular interest as they overlap with miRs that have also been reported as dysregulated in one or more human OA studies (Diaz-Prado et al., 2012; Iliopoulos et al., 2008; Jones et al., 2009; Song et al., 2013).

In the human OA studies, of the 50 dysregulated miRs reported only 6 were common between any one of the studies. In fact, none were common to all 4 studies and only 1 miR (miR-25) was common to 3 of the studies. The striking lack of overlap between the human studies emphasizes the patient and pathological variability of available OA cartilage from clinical samples, which makes interpreting the significance of the human OA-miRs and prioritizing miRs for further functional studies and then therapeutic development difficult. Our studies which interrogate genetically matched samples from a clinically reproducible OA model overcome this problem and provides confidence that our confirmed 7 miRs candidates (Group A) are representative of both early and late-stage OA. Importantly, when we conducted paired analysis of miR dysregulation with the parallel changes in mRNA expression in DMM cartilage (Bateman et al., 2013), all Group A miR candidates showed the corresponding cartilage expression changes of predicted or experimentally determined mRNA targets, compatible with the direction of miR expression change (Fig. 3; Tables 1&2). The expression of all 7 confirmed Group A miR candidates were significantly down-regulated in the protected Jaffa mouse, and therefore, were independent of aggrecan breakdown and early cartilage pathology, suggesting these may be crucial in the mechanisms initiating the pathological cascades that result in eventual cartilage destruction. Thus, miR-15a/16-5p, miR-26b-5p, miR-30c-5p, miR-98-5p, miR-149-5p, miR-210-3p and miR-342-3p offer important potential targets in a critical and wide therapeutic window.

miR-15 and miR-16 are of particular interest as they form a family of related small non-coding RNAs clustered within 0.5kb in the human genome. Both miR-15a-5p and miR-16-5p were down regulated in OA cartilage at both 1 and 6 weeks DMM (Fig. 5A,B). Numerous studies have found miR-16 to be differentially expressed in human OA cartilage and plasma samples (Borgonio Cuadra et al., 2014; Iliopoulos et al., 2008; Lisong et al., 2015; Murata et al., 2010). SMAD3, which has important roles in cartilage development and inflammation, was identified as a target gene of miR-16 in human chondrocytes (Lisong et al., 2015). Furthermore, *Ptgs2* and *Npr3*, both highly up-regulated in OA cartilage, were identified as experimentally validated targets of miR-15/16 seed sequences (Table 2), and both genes are known to have important roles in cartilage homeostasis and OA pathological processes (Bateman et al., 2013; Peake et al., 2014). Collectively, miR-15 and miR-16 may represent high-priority miRs candidates with the potential to contribute to OA initiation and progression. More work is needed to conclusively determine their role in OA pathogenesis.

The remaining high-priority Group A miRs candidates common to human and mouse OA also have targets and functions relevant to OA pathology. Target prediction analysis revealed the up-regulation of *Col10a1* may be mediated by the significant down regulation of miR-26b-5p (Table 1), consistent with the early detection of chondrocyte hypertrophy (Fig. 1). Moreover, the inhibition of miR-26b-5p in IL-1β treated human chondrocytes resulted in the up-regulation of catabolic genes such as MMP-3, -9, -13 and Ptgs2 (Yin et al., 2017). It was particularly striking that *Ptgs2* (Cox2) was identified as a common target between several other dysregulated candidate miRs in our study (miR-98-5p, miR-26b-5p, miR-15a-5p and Let-7d-5p; Tables 1-2; Supplemental Tables 4-5). Ptgs2/Cox2 is involved in the biosynthesis of prostaglandin E2 which is a major catabolic and inflammatory mediator of cartilage degradation. The therapeutic inhibition of Ptgs2 through the use of selective and non-selective inhibitors has been extensively tested in clinical trials for the management of pain and inflammation associated with OA (Laine et al., 2008; Mathew et al., 2011; van Walsem et al., 2015). With varying degrees of effectiveness, and controversy surrounding the safe clinical use of cox-2 inhibitors for the management of OA, the importance of cox-2 inhibition specifically still remains to be determined (Fukai et al., 2012; Katz, 2013; Mathew et al., 2011). Nevertheless, the regulation of *Ptgs2* by several OA-dysregulated miRs identified in this study is of high interest and warrants further pre-clinical studies examining the importance of these miRs to OA pathology and their potential as alternative, more effective therapeutic targets.

Furthermore, several members of the miR-30 family have also been shown to be differentially expressed in human OA (Iliopoulos et al., 2008; Li et al., 2015; Song et al., 2013). Specifically, we demonstrated miR-30c-5p to be significantly down regulated in DMM cartilage, and IPA target analysis revealed *Npr3* and *Nt5e* as experimentally validated targets of miR-30c-5p (Table 1). Npr3 (C type natriuretic peptide) signalling is important in maintaining cartilage function and induces catabolic responses via MEK/ERK signalling and stimulation of anabolic activities via Erk1/2 in an attempt to maintain homeostatic function (Peake et al., 2014). Npr3 not only has interesting OA related activities but it is also potentially targeted by miR-15/-16 and miR-149 suggesting shared regulatory networks between the miRs. *Nt5e* encodes CD73, deficiency of which leads to tissue calcification and early onset OA in hands (Ichikawa et al., 2015).

miR-98-5p has also been linked to OA-relevant pathways and has been shown to have roles in inflammatory processes by modulating IL-1β mediated production of TNFα (Jones et al., 2009). Intriguingly, the down regulation of miR-98-5p has been associated with an increased rate of apoptosis in human chondrocytes and exogenous injection of miR-98 mimic into an OA rat model demonstrated promising therapeutic benefit (Wang et al., 2016a). Conversely, another rat OA study reported contradicting findings of miR-98 over-expression leading to detrimental effects on chondrocyte apoptosis and cartilage integrity (Wang et al., 2016b). More studies are needed to conclusively determine the functional role of miR-98.

The expression of other pro-inflammatory cytokines (TNFα, IL1β and IL6) has been shown to be regulated by miR-149-5p, thus highlighting miR-149-5p as an important inflammatory mediator in OA pathogenesis (Santini et al., 2014). Similarly, miR-210-3p was found to have regulatory roles in inflammation/cytokine expression (Qi et al., 2012), and its over-expression in OA rats decreased inflammation by inhibiting the NF-kB pathway, reducing cytokine production, leading to anti-inflammatory and anti-apoptotic effects (Zhang et al., 2015). Presently, the role of miR-342 in cartilage is unknown. However, our miR-mRNA target prediction analysis identified the appropriate directional dysregulation of several predicted targets in OA mice, including *Bmp7, Inhba* and *Il-11* (Table 2) which are very plausible components of OA networks. Collectively, these studies confirms the validity of our approach for discovery and prioritising these high-priority candidate miRs for comprehensive functional and therapeutic assessment in the future.

Of the remaining 45 miRs following integrated miR-mRNA target analysis that did not overlap with findings from the 4 human end-stage OA studies (termed Group B miR candidates), we confirmed miR-574-5p, miR-31-5p and Let-7d-5p as novel OA-dysregulated miR candidates. No information could be found eluding to Let-7d-5p function in cartilage biology or OA. On the other hand, miR-574 was up-regulated during the early phases of mesenchymal stem cell differentiation towards chondrocytes (Guerit et al., 2013). Importantly, over-expression of miR-574 resulted in inhibition of aggrecan and type II collagen expression, both key pathological hallmarks of OA, via an RXRα-Sox9 feedback mechanism (Guerit et al., 2013). miR-574 has also been shown to target members of the β-catenin/Wnt signalling pathway (Ji et al., 2012), which we know to be pathologically dysregulated in OA (Bateman et al., 2013). miR-31-5p was down-regulated in mesenchymal stem cells (MSC) derived from degraded cartilage compared with MSCs from healthy apparent cartilage from the same donor (Xia et al., 2016). miR-31-5p was also predicted computationally to be a potential regulator of TGFβ3/BMP2-driven processes during chondrogenesis of MSCs (Sang et al., 2014), although thorough experimental investigation is required. Further evidence has demonstrated miR-31 as a negative regulator of osteogenesis of human MSCs by regulating the expression of SATB2, a critical modulator of the expression of osteogenic specific genes (Xie et al., 2014). Nevertheless, the importance of miR-31-5p, miR-574-5p and Let-7d-5p in the context of cartilage/chondrocytes function and OA pathology remains to be explored.

Whilst the role of each individual high-priority miR candidate may have a significant influence on OA molecular processes, miRs function collectively in complex-rich, highly connected miR-mRNA networks (Bracken et al., 2016). Therefore, to gain a more holistic insight into the functional processes significantly impacted by OA-dysregulated miR-mRNA interactions during early OA initiation where we observed the most miR dysregulation (1 week DMM), functional enrichment analysis of miR targets was conducted (Fig. 8). Significant biological processes (adjusted p. value < 0.05) disrupted during early OA were identified using ToppGene (Chen et al., 2009) and then clustered using REViGO (Supek et al., 2011) (Fig. 8). Functional annotation of the putative miR dysregulated mRNA targets revealed processes largely involved with regulation of MAPK cascade, positive regulation of signal transduction, cell migration and development (Fig. 8). ECM organization, response to mechanical stimulus and regulation of cell adhesion were also clusters significantly affected by dysregulated miR targets (Fig. 8). These OA-relevant cellular processes, significantly disrupted during early OA initiation (1 week DMM) but prior to OA pathology, provides confidence that our high-priority miRs are exciting upstream candidates for therapeutic intervention and targeting.

**Figure 8.**
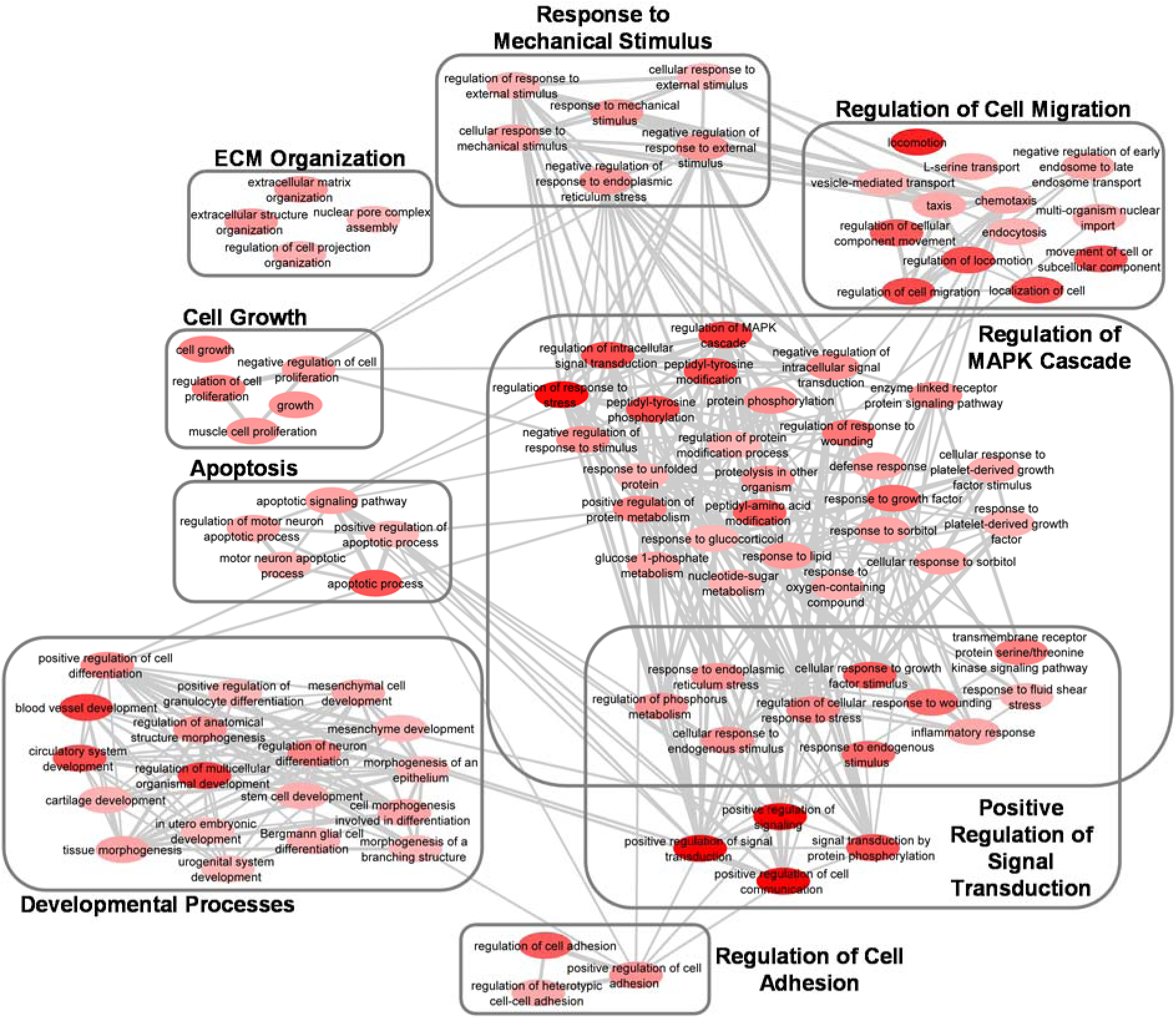
Gene Ontology (GO) biological processes associated with dysregulated miR targets during early OA initiation. Functional enrichment analysis using ToppGene was performed on putative miR targets to highlight biological processes most significantly affected by dysregulated miR-mRNA interactions during early OA initiation. GO terms (adjusted p. value < 0.05) were summarized and visualised using REViGO and Cytoscape. Boxes represent the top clusters of biological processes significantly influenced by dysregulated miR targets. Node colour intensity signifies adjusted p. value, with highly significant GO terms in dark red. Highly similar GO terms are linked by edges, where the line width indicates the degree of similarity.

We profiled miR expression in the adjoining sclerotic SCB tissue as it is well known that cartilage and SCB have a dynamic relationship involving biochemical and molecular crosstalk between the two tissues (Sharma et al., 2013; Yuan et al., 2014). However, we were unable to detect statistically significant dysregulation of miRs in DMM versus sham SCB tissue. Although ~ 80% of the total number of miRs expressed and dysregulated in cartilage were also found in SCB tissue, none of these were significantly differentially regulated in OA SCB. Collectively, this supports the notion that OA is associated with cartilage-specific regulation of common miRs rather than the regulation of cartilage-specific miRs.

The lack of differential expression in SCB samples was not due to lack of sensitivity of our profiling approach as statistically significant temporal changes in miR expression were evident. This clearly demonstrates the sensitivity of the microarrays to detect miR expression changes if present, highlighting the robustness of the dataset. In contrast with our findings, two studies have previously identified miR dysregulation in SCB tissue from human end-stage OA (Jones et al., 2009; Prasadam et al., 2016). Both demonstrated differential expression of 30 miRs in human OA bone tissue isolated during total knee replacement operations. The obvious discrepancies between our studies (species, model, stage of disease progression) may account for the lack of commonality between our datasets. Patients undergoing total knee surgery represent end-stage OA and exhibit substantial SCB changes impacted by years of progressive OA. In comparison our studies were conducted over a 6 week time period and thus recapitulate a mild early pathology. Investigating later time points after DMM would confirm whether miR changes occur with advanced pathology, however therapeutic intervention at such late stage of disease would be limited from a disease modifying prospective.

While our data suggest that miR dysregulation in the SCB is not a significant component of the pathophysiology of OA initiation and early progression in this model, it is important to acknowledge unavoidable technical limitations which may mitigate against detecting SCB miR dysregulations. Despite using a laser micro-dissection approach to isolate RNA from the localised regions of SCB it was impossible to exclude some regions of the surrounding bone marrow. As a result, any SCB-specific pathological changes may be masked by inclusion of RNA from bone marrow. Nonetheless,

For the first time, we have described potential cartilage miR regulators of early OA initiation and progression that share common findings with end-stage human OA data and also novel miRs which have not been previously associated with OA. *In silico* and experimental data generated target prediction algorithms on paired cartilage miR:mRNA expression, identified interesting miR candidates with targets implicated OA pathology. Importantly, our studies provide critical information on the role and biological processes of cartilage miRs during the early stages of OA initiation and progression not possible with patient tissues. Studying the early stages of the disease is pivotal mechanistically and because it represents a critical time when therapeutic intervention is expected to have the most disease modifying outcome. Our work provides a robust platform and validated pool of high-priority OA-associated cartilage miRs. Further studies are underway to determine the role of specific individual miRs and their potential as therapeutic targets in OA disease initiation and progression.

## MATERIALS AND METHODS

### Animal models and induction of OA

The generation of mice resistant to aggrecanase cleavage in the aggrecan IGD (Jaffa mouse; *Acan* p.374ALGS→374NVYS) has been previously described (Little et al., 2007). Post-traumatic OA was surgically induced in 10-12 week old male wild type C57BL6 and Jaffa mice by bilateral destabilization of the medial meniscus (DMM), as described previously (Glasson et al., 2007; Jackson et al., 2014). Briefly, under anaesthesia the medial menisco-tibial ligament was exposed (by medial *para*-patellar arthrotomy and intrapatellar fat pad elevation, without tissue resection) and transected with curved dissecting forceps by one surgeon (CBL). Bilateral sham-operations were also performed, where the medial menisco-tibial ligament was visualized but not transected. All joints were flushed with sterile saline to remove any blood prior to separate closure of the joint capsule (simple continuous 8/0 polyglactin 910), subcutaneous tissue (mattress suture 8/0 polyglactin 910) and skin (cyanoacrylate).

DMM and sham mice were co-housed with 2-5 animals/30×20×18cm individually-ventilated-cage with filter lids, provided with sterilized bedding and environmental enrichment, maintained at 21-22°C with a 12-hour light/dark cycle, and received water and complete pelleted food *ad libitum*. Mice received no post-operative medication, were maintained in their pre-operative groups and were allowed unrestricted cage exercise.

Mice were randomly allocated to treatment groups/harvest time point prior to study commencement using their individual ID numbers. Animals were sacrificed at 1 and 6 weeks after surgery. For each animal, one joint was processed for histology whilst the other was used for microarray expression profiling/qPCR. Microarray experiments on cartilage samples were performed on n = 3/group/time point, each consisting of RNA pooled from 3 individual mice. qPCR validation was performed on the same 3 pooled RNA samples and 4 additional biological replicates per time point per group. Microarray experiments on SCB samples were performed on n = 4 mice/group/time point with qPCR validation performed on a different cohort of 4 mice/group/time point.

### Histopathological analysis of OA features

Knee joints were dissected, fixed, decalcified, processed for histology and scored for OA histopathologic features as described previously (Jackson et al., 2014). Sagittal sections every 80 μm across the medial femoro-tibial joint were collected and stained with toluidine blue and fast green. Cumulative scores (total in all sections) of cartilage and SCB OA histopathologic features, which were determined by two independent observers who were blinded with regard to surgical intervention and post-operative time, were averaged, and these mean values provided a single histologic score for each of the following features in each mouse: cartilage proteoglycan loss (score range 0-3), structural damage (score range 0-7), chondrocyte hypertrophy (score range 0-1), SCB sclerosis (score range 0-3), osteophyte size (score range 0-3) and maturity (score range 0-3). Only histologic scores for the tibia are presented since miR profiling was performed on tibial articular cartilage only and can be correlated with adjacent SCB pathologic changes.

### Laser capture micro-dissection and RNA preparation

Joint dissection, laser micro-dissection and RNA extraction from mouse cartilage and SCB was performed in a similar fashion to previously described (Bateman et al., 2013; Belluoccio et al., 2013). Briefly, mice were sacrificed 1 and 6 weeks after surgery and the tibial epiphyses were isolated and placed in RNA Later (Life Technologies) containing 10% EDTA pH 5.2 for at least 72 hrs at 4°C with gentle agitation. After decalcification the samples were washed in DEPC-treated PBS, embedded in OCT and stored at -80°C. Serial 10 μm sagittal cryo-sections on polyethylene naphthalate membrane-coated slides (Thermo Fisher Scientific) were fixed in ethanol and air-dried. Sections of the cartilage and underlying SCB of the medial tibial plateau were laser micro-dissected (Arcturus Bioscience) and collected. Total RNA was extracted from pooled laser micro-dissected sections from each individual mouse joint using TRIzol according to the manufacturer’s instructions. RNA integrity and quantification was determined by capillary electrophoresis on a TapeStation 2200 with a high sensitivity screentape (Agilent Technologies) according to the manufacturer’s specifications.

### miR expression profiling

miR expression profiling was performed using SurePrint mouse miR microarray technology, release 21 (G4859C, Agilent Technologies) at the Ramaciotti Centre for Genomics (UNSW, Sydney, Australia). Briefly, 70 ng and 100 ng of total RNA from cartilage and SCB, respectively, was labelled and hybridized using the miR microarray System with miR complete labelling and Hyb kit version 3.0 (Agilent Technologies) by following manufacturer’s instructions. The arrays were scanned on a G2565CA microarray scanner and the features were extracted using Agilent Feature Extraction 12.0.07 software.

### Integrated miR-mRNA analysis and functional enrichment analysis

Bioinformatic analysis to identify putative miR targets was performed by uploading miR:mRNA expression data (Bateman et al., 2013) into the MicroRNA Target Filter module within Ingenuity Pathway Analysis (IPA, Qiagen Redwood City, CA, USA) application to generate a network of putative miR gene targets directly relevant to OA cartilage. Inversely expressed miR-mRNA interactions were selected and functional enrichment analysis of miR targets was performed using ToppGene (Chen et al., 2009) with Benjamini-Hochberg false discovery rate adjusted p-value cut-off < 0.05. Gene Ontology (GO) terms and associated p-values generated through ToppGene were then summarized and visualised using REViGO (Supek et al., 2011). Interactive graphs were subsequently generated using Cytoscape software (Shannon et al., 2003).

### Quantitative polymerase chain reaction (qPCR)

qPCR validation was carried out on a LightCycler 480 Instrument (Roche) using the miScript microRNA RT qPCR system (Qiagen) for miR expression by following manufacturer’s instructions. All qPCR reactions were performed in duplicate and U6 snRNA and SNORD 61 snRNA were used as internal reference controls.

### Statistical analysis

Comparison of histopathology scores between treatment groups and time points were analysed using nonparametric ranked Kruskal-Wallis analysis for multiple groups, and where there was significance (p<0.05) the Mann Whitney U-test (for unpaired data) was performed for between group comparisons with Benjamini-Hochberg correction for multiple comparisons (StatSE software, Stata corporation, TX, USA).

The microarray data was background corrected using the Normexp method and normalized with cyclic loess as previously described (Kung et al., 2017) using the bioconductor package Limma (Limma_3.20.9; (Ritchie et al., 2015)). Only probes with 10% greater signal than the negative controls in at least 4 samples were maintained for differential expression analysis. Probes were summarized initially at the probe level then at the gene level using the Limma avereps function, with control probes removed. Differential expression was assessed using moderated t-tests from the Limma package. Results were adjusted for multiple testing using Benjamini-Hochberg method to control for false discovery rate. The data has been submitted to the GEO data repository (https://www.ncbi.nlm.nih.gov/geo/; accession number GSE93008 (SCB); accession number for cartilage in progress).

miR qPCR expression data was quantitated using the comparative Ct method and the results were analyzed by nonparametric unpaired Mann-Whitney U-test to evaluate differences between group comparisons (GraphPad Prism, version 7.01).

### Study approval

All animal procedures were approved by the Royal North Shore Hospital Animal Ethics Committee (Protocol RESP/14/77).

## AUTHOR CONTRIBUTIONS

Conception and design: JFB, CBL

Collection and assembly of data: VR, LHK, LR, CA

Analysis and interpretation of the data: LHK, KMB, JFB, CBL

Providing reagents: AJF

Drafting of the article: LHK, KMB, JFB, CBL

Critical revision of the article for important intellectual content: All authors

Final approval of the article: All authors

Obtaining of funding: JFB, CBL

## ACKNOWLEDGMENTS

Supported by the National Health and Medical Research Council of Australia Project Grant APP1063133 and by the Victorian Government’s Operational Infrastructure Support Program. The authors thank Ms Susan Smith for processing and sectioning the mouse knee joints, and blinding and randomising sections for scoring.

